# An improved rapid and sensitive long amplicon method for nanopore-based RSV whole genome sequencing

**DOI:** 10.1101/2025.01.06.631406

**Authors:** Xiaomin Dong, Steven Edwards, Yi-Mo Deng, Clyde Dapat, Arada Hirankitti, Rachel Wordsworth, Paul Whitney, Rob Baird, Kevin Freeman, Andrew J Daley, Ian G. Barr

## Abstract

**Background:** Whole-genome sequencing (WGS) provides critical insights into the Respiratory syncytial virus (RSV) transmission and any emerging mutations that could impair the efficacy of monoclonal antibodies or vaccines that have been recently licenced for clinical use worldwide. However, the ability to sequence RSV genomes at large scale is limited by expensive and time-consuming sequencing methods. Oxford Nanopore Technology (ONT) offers significant improvements in next generation sequencing (NGS) both in turnaround time and cost, compared to other platforms for viral WGS.

**Methods:** We have developed and modified an RSV long amplicon-based WGS protocol for the ONT platform using a one-step multiplex RT-PCR assay and the rapid library barcoding kit. 135 RSV positive Australian clinical specimens (91 RSV-A and 44 RSV-B) sampled in 2023 with Cycle threshold (Ct) values between 14 to 35 were tested in this study. This ONT workflow was compared with other recent RSV WGS amplification assays based on short amplicons.

**Results:** A PCR amplicon clean-up step prior to library preparation significantly improved WGS result for samples with poor amplicon generation, but it is not necessary or beneficial for ones that generated high concentrations of amplicons. Overall, a success rate of 85.9% was achieved for WGS. This method performed as well as the more complex short amplicon methods in terms of genome coverage and sequencing depth.

**Conclusions:** The workflow described here was highly successful in generating RSV WGS on ONT platform and had improved turnaround times and excellent results with RSV clinical samples with Ct values up to 30.

## 1. Introduction

Human Respiratory Syncytial Virus (RSV) is a respiratory pathogen that can have a severe impact on human health, especially in infants, young children, older adults, and immunocompromised individuals (1, 2). RSV is a single-stranded negative-sense RNA virus with a genome of 15.2 kb in length that can be transcribed into at least 11 proteins.

Recently, a long-acting monoclonal antibody (Mab) (Nirsevimab) and three RSV vaccines have been introduced in the market to reduce the burden of RSV in newborns and in people aged 60 years and over (3-8). All of these RSV preventatives target the RSV pre-fusion (F) protein, a key surface protein that is responsible for fusing the virus and cell membranes and allowing release of the viral genome into the cytoplasm and subsequent virus replication(9). Two of these vaccines are based on recombinant pre-F proteins (one for the pregnant women and persons over 60 years of age (Abrysvo^™^) and another for those over 60 years of age (Arexvy^™^)) and one based on mRNA which encodes the pre-F protein (for persons over 60 years of age (mRESVIA^™^)). In addition, other Mabs and vaccines to combat RSV that also target the F protein or other viral proteins such as L or NS1 protein are in various stages of clinical development as well as live attenuated RSV vaccines (5). Therefore, ongoing surveillance of circulating RSV strains by sequencing of the entire viral genome is needed to monitor viral evolution and the rise and spread of mutations that might generate resistance to these newly introduced preventatives, that could reduce their effectiveness against circulating RSV. The loss in effectiveness of several Mabs Antibodies against viral infections due to viral evolution was exemplified during the recent SARS-CoV-2 pandemic (10) as well as an RSV Mab which failed due to a 2–amino acid substitution in the RSV-B in the Suptavumab^™^ epitope that led to loss of neutralization activity (11).

Next generation sequencing (NGS) has become an invaluable tool in genomic surveillance of infectious diseases. It has greatly facilitated the monitoring of SARS-CoV-2 evolution, especially the appearance of concerning variants with enhanced infectivity and transmissibility during COVID-19 pandemic period (12). However, obtaining full genome coverage with NGS can be challenging as it requires an even coverage with high sequencing depth to accurately detect important mutations and polymorphisms. Amplicon-based NGS assays have been widely used for viral whole-genome sequencing as they are generally sensitive, cost- and time-effective methods. With the SARS-CoV-2 pandemic, primers designed with the Primal Scheme pipeline that generated tiled PCR fragments fully covering viral genome (13), known as the real-time molecular epidemiology for outbreak response (ARTIC) protocol, were popularised for sequencing either on the Illumina or Oxford Nanopore Technologies (ONT) platform (13-15). This protocol was developed by the ARTIC network based on an earlier strategy for sequencing single-stranded RNA viruses from high cycle threshold (Ct) clinical samples and involved the use of nearly 100 primer pairs to cover the ∼ 30kb genome of SARS-CoV-2 (16). When comparing sequencing platform, ONT sequencing has the advantages of real-time read-out of sequencing, long read length, portability, short turnaround time and low cost compared to the Illumina platform, making it highly suited to rapid viral pathogen detection and whole-genome sequencing (WGS), as was seen in outbreak investigations of SARS-CoV-2 (17).

For RSV, sequencing the smaller full-length genome rather than partial genome covering the *G* and/or *F* genes provides more information and is therefore a more powerful tool for studying virus evolution and identifying mutations associated with current and future interventions. We previously established a long amplicon method for RSV based on one-step multiplex RT-PCR (mRT-PCR) followed by NGS on either Illumina or ONT platform for RSV WGS (18) using only two PCR reactions per sample. Here, we describe an improved method using the ONT NGS workflow and the ONT Rapid barcoding kit (RBK) which enabled easier library preparation with PCR amplicons resulting in a simple, rapid, and cost-effective method for RSV whole-genome sequencing (WGS). This method was compared to other “ARTIC-like” methods that have been developed recently, in terms of the NGS coverage.

## 2. Methods and materials

### 2.1 Clinical specimens

De-identified RSV positive respiratory samples were used in this study. Sample types including nasal swabs, nasopharyngeal swabs, nasal washes, or nasopharyngeal aspirates were collected between September and November 2023 from Australian patients aged from 0 to 84. Samples were shipped to the Centre and stored at -80°C until analysis.

### 2.2 RNA extraction and real-time RT-PCR assay for characterizing RSV subgroups and viral loads

QIAamp 96 Virus QIAcube HT (QIAGEN) reagents were used to extract viral RNA from 200 μl of clinical samples according to manufacturer’s instruction. RSV subgroups (RSV A or B or mixed) and viral loads were determined with the CDC Respiratory Syncytial Virus Real-time RT-PCR panel (RSV_RUO-01) as described previously (18, 19). The resulting cycle threshold (Ct) values for evaluating viral load in samples were rounded to the nearest whole number in this study.

### 2.3 RSV amplicon generation by one-step mRT-PCR for RSV WGS

SuperScript IV One-Step RT-PCR System with ezDNase (Invitrogen) was used for the long PCR amplicon protocol (Supplementary Table 1). Primers and RT-PCR conditions were modified to accommodate some changes in the more recent RSV sequences (18). Briefly, DNase treatment of extracted RNA was performed according to the manufacturer’s instructions for human genomic DNA removal. The reactions were set up according to Supplementary Table 2. Thermocycling conditions were 10 (minutes) min at 50°C for reverse transcription, 2 min at 98°C for reverse transcriptase enzyme inactivation followed by 40 cycles of 98°C for 10 seconds (s), 55°C for 30 s and 72°C for 3 min, and final extension step at 72°C for 5 min. mRT-PCR products were quantified with QuantiFluor dsDNA System reagents (Promega) according to the manufacturer’s instructions and read on a FLUOstar Omega Microplate Reader (BMG Labtech). mRT-PCR products from representative samples were also quantitated on TapeStation using D5000 Tapes for visualization. An equimolar mixture of RT-PCR products from the two reaction tubes per sample were normalized and used for downstream ONT NGS library preparation and sequencing.

Two ARTIC-like short amplicon WGS methods were also compared to the long amplicon method in this study. The first protocol was developed by Maloney *et al* and modified by us with Invitrogen SuperScript IV One-Step RT-PCR System and similar thermocycling conditions as described above for the long PCR amplicon assay for amplicon generation except with 72°C for 30 s instead of 3 min (20). A second ARTIC-like protocol developed by Talts *et al* was carried out according to their published method (21).

### 2.4 ONT library preparation and sequencing on an ONT platform

ONT NGS libraries were prepared with ONT Rapid Barcoding Kit (SQK-RBK114.96) or Rapid PCR Barcoding Kit (SQK-RPB114.24) according to the manufacturer’s instructions. Briefly, NGS libraries were prepared with Rapid Barcoding Kit by using 50 ng of normalized amplicons supplemented with water to 10 μl or Rapid PCR Barcoding Kit by using 2 ng of normalized amplicons supplemented with water to 3 μl as input. PCR product clean-up was performed with AMPure XP Beads (Beckman Coulter) supplied in the kits or similar DNA binding beads with 1:1 of beads-to-sample ratio. PCR product clean-up and enrichment was carried out the same as above steps but with all normalized PCR products used for purification and eluted in 10 μl EB buffer from ONT RBK kit.

### 2.5 Data Analysis

Consensus sequences submitted to GISAID (https://gisaid.org/) were generated using in-house bioinformatic pipeline modified based on IRMA pipeline (18, 22). Version 1.5.1 ARTIC pipeline (https://github.com/artic-network/fieldbioinformatics) was used to better trim off PCR primers and evaluate coverage breadth and depth for all three assays tested in this study. For coverage breadth evaluation, genome coverage depth was set at a minimum read depth of 40x.

## 3. Results

### 3.1 The performance of ONT rapid barcoding method for RSV WGS

A total of 91 RSV-A and 44 RSV-B, with Ct values ranging from 14 to 35, were tested (Supplementary Figure 1A). All six amplicons with similar yields were generated from representative RSV-A and RSV-B samples as visualized by TapeStation analysis (Supplementary Figure 1B).

Samples were divided into two groups based on the quantity of amplicons generated: 115 samples with sufficient long amplicons (≥50 ng of normalized amplicons in 10 µL) were chosen for a standard library preparation with the ONT Rapid barcoding kit (RBK), and 20 samples that had insufficient amplicons (with less than 50 ng of normalized amplicons in 10 µL) were selected for the low input protocol (Figure 1A).

From the 115 samples with sufficient amplicons, RSV WGS were obtained from 89 samples with Ct values up to 28 (77.4%), while incomplete RSV genome sequences but with full-length *G* and *F* genes (GF) were obtained from a further 4 viruses, 4 samples had partial RSV genome with complete *G* gene (G) (Figure 1B). As shown in Figure 1C, both representative RSV-A and RSV-B samples had complete coverage and a minimum depth above 40 reads throughout the whole genome.

Next, we tested if PCR amplicon clean up prior to library preparation could further improve WGS results as this step is recommended in the Rapid Barcoding kit manual from ONT for the removal of PCR artifacts. Twenty-six samples which failed to generate full length sequences were chosen for PCR amplicon clean up. Most of them had low concentrations of amplicons. After PCR clean up, 13 out of 26 samples generated complete genomes (Figure 1D). Furthermore, we did a side-by-side comparison on 5 samples, with the same PCR products divided equally into two parts, one for library preparation directly and one for amplicon purification followed by library preparation. The resulting libraries were loaded on the same flow cell ONT run for NGS data output comparison. More RSV specific NGS reads (about 13-fold increase) was observed from libraries made from purified amplicons compared to non-purified amplicons for both RSV-A and RSV-B samples for these 5 samples (Figure 1E-G). However, the same amplicon purification step was also tested on 30 samples with high concentration of amplicons (>20ng/µL normalised amplicons), and results showed no improvement. (Figure 1H).

**Figure 1.**
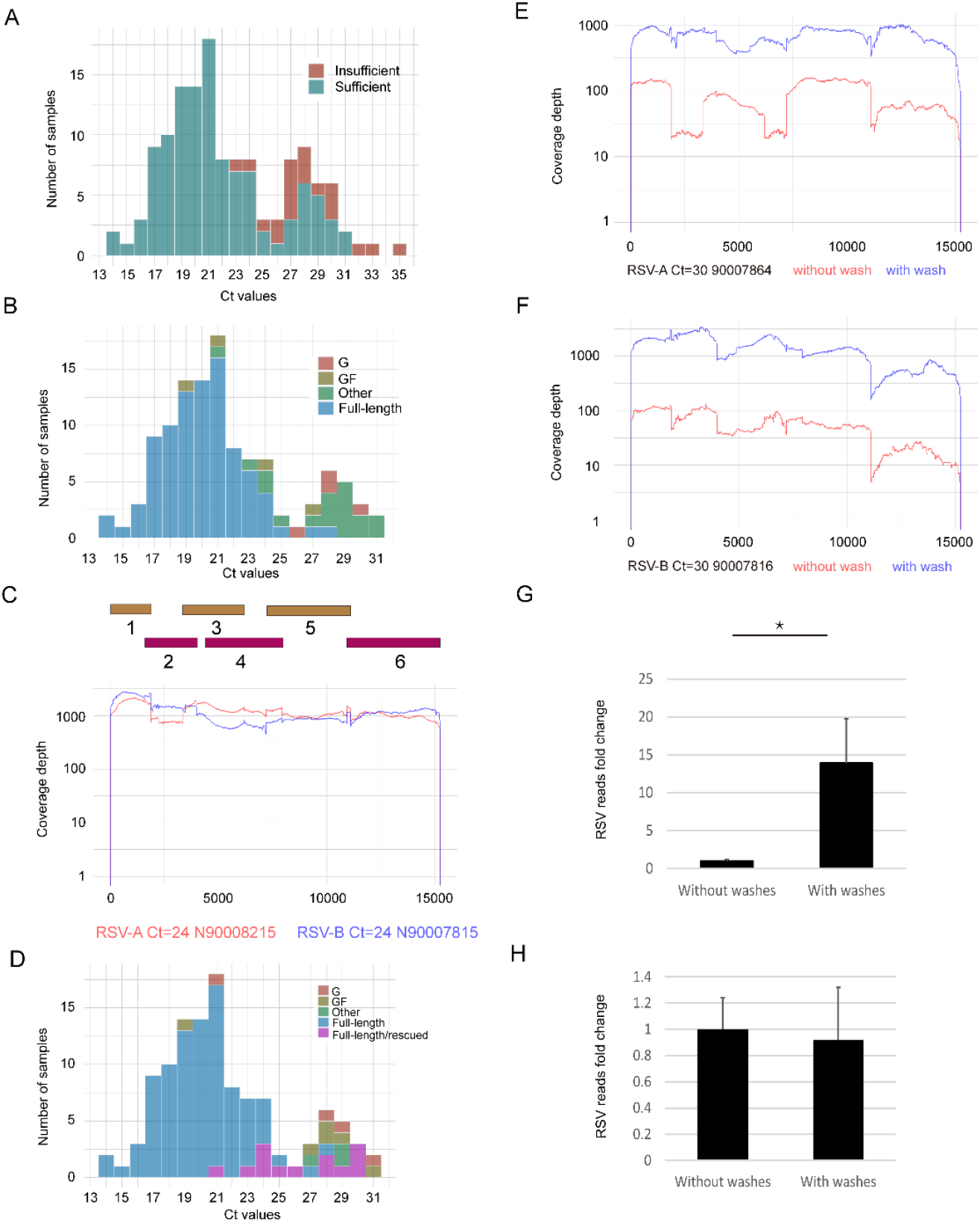
The performance of ONT rapid barcoding method mediated NGS with library preparation from samples with sufficient amplicons. (A) Histograms of the Ct value distribution for clinical samples having sufficient (dark green) and insufficient (dark red) amplicons generated for ONT libraries. (B) Histograms of the Ct value distribution for clinical samples within sufficient amplicons group with RSV full-length genome (blue), *G* plus *F* genes (brown), only G gene (dark red) and missing *G* and *F* gene sequenced (also called “other”), respectively. (C) Coverage depth of sequenced representative RSV-A (red) and RSV-B (blue) in genomic position covered by six overlapping amplicons. (D) Histograms of the Ct value distribution for clinical samples within sufficient amplicons group having RSV whole genome (blue), *G* plus *F* genes (brown), only G gene (dark red) and missing *G* and *F* gene sequenced (also called “other” in dark green), whole genome rescued by DNA purification (purple), respectively. Coverage depth of sequenced representative RSV-A (E) and RSV-B (F) in genomic position with (blue) and without (red) PCR amplicon clean-up. Bar plots showing the fold change of NGS reads mapped to RSV reference genomes from sequenced samples of low (G) and high (H) concentration of amplicons generated without and with PCR amplicon purification. NGS reads of libraries from washed PCR amplicons were normalised and expressed as fold changes to ones from unwashed PCR amplicons that were set to 1. Data are expressed as mean ± SD. *t-test* analysis was performed for statistical significance. *P* values less than 0.05 were considered as statistically significant and labelled as * in the figures.

### 3.2 Comparison of sequencing samples with insufficient amplicons for ONT NGS library preparation by using rapid barcoding (RBK) and rapid PCR barcoding (RPB) kits

To further increase the sensitivity of ONT sequencing, we investigated two protocols for samples with very low quantity of PCR products produced (normalised amplicons < 5 ng/µL): one was to add a purification and concentration step with DNA binding beads before RBK library preparation; the other was to use the rapid PCR barcoding (RPB) kit which included 20 PCR cycles to amplify the library before loading it to flow cells. Twenty samples with insufficient amplicons (normalised amplicons < 5 ng/µL) were used for this comparison. Better coverage and depth were achieved using the ONT RBK kit with a simple concentrating step compared to RPB kit (Supplementary Figures 2A and 2B, Supplementary Table 3).

### 3.3 Comparison with ARTIC-like methods for RSV WGS

Our method was compared with two short PCR amplicon ARTIC-like based NGS, one with an average amplicon size of 400 bp by Maloney *et al* (hereinafter referred to as Maloney’s method) and the other one with amplicons ranging from 630 bp to 1000 by Talts *et al* (hereinafter referred to as Talts’ method) (21). Ten RSV-A samples with Ct’s between 22 and 33 and ten RSV-B samples with Ct values from 21 to 30 were chosen for this comparison. As shown in Figure 2, good and even coverage depth was achieved by Talts’s method for both representative RSV-A and RSV-B samples, which was comparable to that obtained from our assay. However, the coverage depth of Maloney’s method was very patchy and with many gaps across the RSV genome for both RSV-A and RSV-B samples. Overall, the long amplicon method and the Talts protocols successfully generated 10/20 (50%) and 9/20 (45%) WGS from the clinical samples respectively while the Maloney protocol generated 0/20 WGS (Table 1). Moreover, the length of all RSV-A genomes sequenced with both short amplicon-based PCR assays were slightly shorter than those generated with our assay (Figure 2A). All ten tested RSV-A samples had only partial RSV genomes sequenced with both short amplicon generation assays, due to the first 3’ end forward primers being located in RSV-A *NS1* open reading frame (ORF) for both short amplicon assays leading to the loss of 12 nucleotides from the RSV full-length genome. In addition, the Talts’s method was 100 nucleotides short as the final 5’ end reverse primer was located within the RSV-A *L* gene body. PCR amplicon clean-up was performed for samples which failed to generate full length sequences but didn’t improve RSV genome coverage showed in Table 1.

The Talts’ method was however more sensitive than our assay for four RSV-B samples (90008172, 90008161, 90008207 and 90007819) and two RSV-A samples (90008229 and 90008224) in terms of RSV genome coverage (Table 1). DNase treatment of RNA samples was not performed for both Talts’ and Maloney’s methods, but they produced very low human genome background if any, (except for samples of high Ct values such samples 90008144 with Ct at 29, 90008203 with Ct at 33, and 90008224 with Ct at 30), suggesting DNase treatment of RNA prior to the generation of short amplicons is generally not required unlike the long amplification method which is generally improved by DNase treatment.

**Figure 2.**
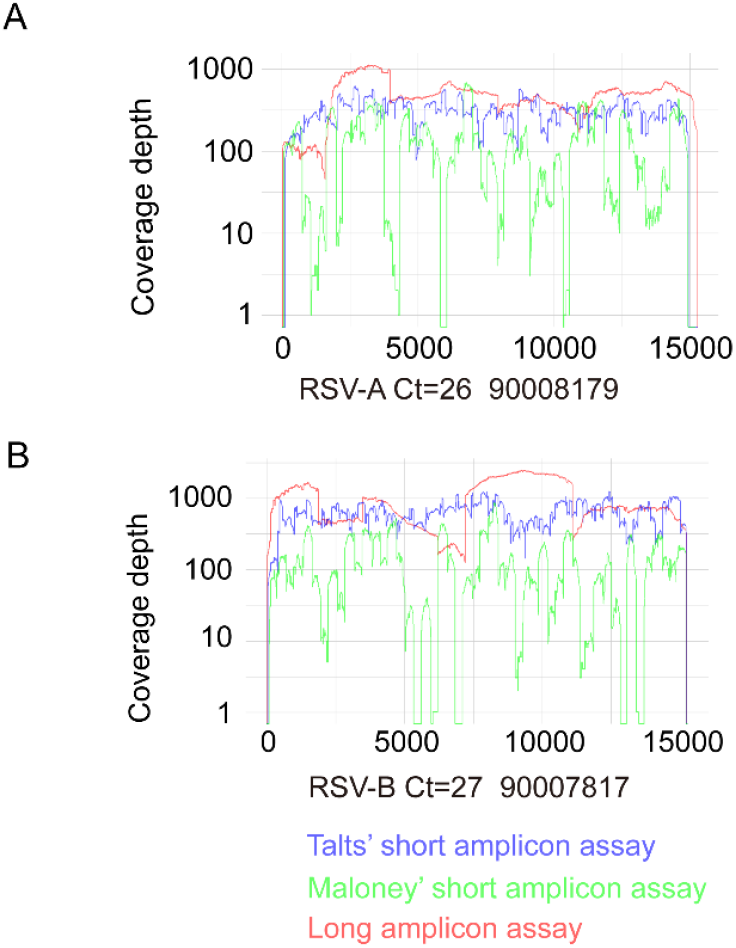
RSV NGS achieved by one long and two short amplicon-based RSV WGS assays. Coverage depth of sequenced representative RSV-A (A) and RSV-B (B) in genomic position with long amplicon-based (red), Maloney’ short amplicon-based (green) and Talts’ short amplicon-based (blue) assays.

**Table 1.** Comparison of long and short amplicon based NGS assays. NGS results were characterised into four groups including Full-length (RSV whole genome sequence obtained), G (Full G sequence only), F (Full F sequence only), GF (Full F and G sequence obtained) and other (no full G and F sequences obtained).

| Sam<br>ple<br>ID | Subt<br>ype | RSV Ct | Coverage of<br>long PCR<br>amplicon<br>WGS | Coverage<br>of Talts'<br>WGS | Human<br>reads for<br>Talts' WGS | Coverage of<br>Maloney's<br>WGS | Human<br>reads for<br>Maloney's<br>WGS |
| --- | --- | --- | --- | --- | --- | --- | --- |
| 9000<br>8203 | A | 33 | Other | Other | 22% | Other | 5% |
| 9000<br>8224 | A | 30 | G | GF | 24% | Other | 17% |
| 9000<br>8235 | A | 31 | G | F | 2% | Other | 0.9% |
| 9000<br>8229 | A | 32 | G | GF | 11% | Other | 11% |
| 9000<br>8197 | A | 28 | G | F | 0.8% | Other | 0.2% |
| 9000<br>8179 | A | 26 | Full-length | GF | 9% | G | 3% |
| 9000<br>8173 | A | 22 | Full-length | GF | 6% | G | 4% |
| 9000<br>8215 | A | 24 | Full-length | GF | 5% | G | 0.5% |
| 9000<br>8234 | A | 23 | Full-length | GF | 1% | G | 0.1% |
| 9000<br>8191 | A | 27 | Full-length | GF | 0.9% | G | 0.06% |
| 9000<br>8144 | B | 29 | Other | Other | 38% | Other | 24% |
| 9000<br>8172 | B | 29 | G | Full-length | 2% | Other | 2% |
| 9000<br>7815 | B | 24 | Full-length | Full-length | 0.5% | Other | 0.3% |
| 9000<br>8161 | B | 28 | GF | Full-length | 8% | Other | 8% |
| 9000<br>8183 | B | 23 | Full-length | Full-length | 0.7% | Other | 0.6% |
| 9000<br>8207 | B | 21 | G | Full-length | 2% | Other | 0.7% |
| 9000<br>8205 | B | 25 | Full-length | Full-length | 3% | Other | 2% |
| 9000<br>7819 | B | 30 | Other | Full-length | 2% | Other | 1% |
| 9000<br>7808 | B | 22 | Full-length | Full-length | 0.3% | Other | 0.2% |
| 9000<br>7817 | B | 27 | Full-length | Full-length | 0.3% | Other | 0.09% |

### 3.4 Phylogenetic and F gene analysis of Australian RSV A and B viruses

All WGS obtained in this study were analysed phylogenetically and showed that the RSV-A genomes clustered into 3 main clades including A.D.1, A.D.3 and A.D.5 (Supplementary Figure 3) while the RSV-B genomes clustered into 2 main clades including B.D.E.1 and B.D.E.4 (Supplementary Figure 4).

Amino acids encoded by the RSV genomes obtained in this study were examined for mutations in the binding sites of the licensed monoclonal antibodies Palivizumab or Nirsevimab. No mutations were found in the Palivizumab-binding site between 256 and 276 amino acids of F protein for both RSV-A and RSV-B sequences (23). In the case of Nirsevimab, 19 out of 38 RSV-B sequences had S211N substitution in F protein, which has been reported previously but does not cause increased resistance to Nirsevimab based on neutralization tests (24).

## 4. Discussion

In this study, we improved our previously published one-step multiplex RT-PCR and ONT workflow using a much simpler and quicker solution for library preparation. With this workflow, a total of 116 RSV WGS were successfully generated from 135 RSV positive clinical samples (85.9%) when setting a minimum read depth of 40x. The portable MinION sequencer provides field sequencing and is already commonly used in sequencing facilities for performing viral whole-genome sequencing, especially for SARS CoV-2. Two library preparation methods are commonly used for amplicon based NGS with the ONT platform. One is ligation-based which was used in the ARTIC protocols and our previous paper, and another that is based on fragmentation was used in this current study and produced as equally good results but was much quicker and simpler.

Several limitations of previously developed multiplex RT-PCR amplification and NGS assays for RSV WGS have been addressed with this updated method. A significant increase in the sensitivity achieved on samples with low RSV viral load, in addition the library preparation time has been reduced along with the cost. Modifications made to our previous method (18) included using the ONT rapid barcoding kit which replaced both the Illumina DNA preparation kit and the ONT ligation sequencing kit (Supplementary Table 1). For samples with lower quantity of PCR amplicons, an extra PCR amplicon clean-up step prior to ONT RBK library preparation significantly improved generation of WGS. This modification worked better than ONT RPB kit for low amplicon input samples in this study with more even coverage depth, probably due to the extra PCR step involved in ONT RPB method (25). However, this PCR amplicon clean-up step is not beneficial for samples that generate higher concentrations of amplicons.

The current workflow was also compared with two short amplicon-based RT-PCR (ARTIC-like) assays for RSV WGS. All RSV-A genomes produced by both short PCR amplicons-based methods had slightly abridged WGS with a 12-nucleotide gap at start of the *NS1* gene due to the primer design. While this small region is not an important site in the RSV genome, it would be preferable for their primers to be modified to fully cover the genome of RSV in the future. The Talts’ method also generated WGS that lacked the final 100 nucleotides at the final 5’ end of the genome, which is within the RSV-A *L* gene body.

Long amplicon and short amplicon based sequencing methods have their own pros and cons. Long amplicon-based multiplex assays only require a small number of primers in each reaction and minimise the chance of introducing too many PCR products amplified differentially in the same tube (26). Independent studies on SARS-CoV-2 WGS demonstrated that long amplicons-based sequencing not only reduced the possibility of amplicon dropout and coverage bias, but also improved overall quality of the consensus sequences (27, 28). In addition, as the mutation rate is normally high in RNA virus, the larger number of primers used in the amplicon generation method for WGS the higher chances of primer mismatch or drop-out it will have. This can contribute to uneven coverage of the WGS and may even cause the loss of a whole region of the genome. Significant loss of SARS-CoV-2 genome coverage was reported due to constant viral evolution and especially short amplicon-assays designed with Primal Scheme (27, 29). Therefore, continuous implementation of new primer sets has been used for SARS-CoV-2 surveillance work to restore WGS capability over the entire COVID-19 pandemic period (29, 30). Moreover, it has been widely reported that multiplex RT-PCR assays were likely to generate uneven amplification across the genome and optimization of the primer concentration was considered as a useful strategy to increase the PCR efficiency in poorly covered regions (31). With a smaller number of primers used for amplifying RSV whole genomes, it is easier for long amplicon-based assays than short amplicon-based ones to update the primers when it is needed to cover new variants for each epidemiological season and optimise RT-PCR reaction to have amplicons generated at similar efficiency (29, 30, 32).

On the other hand, one of the major drawbacks of the long amplicon-based assays is its lower sensitivity for degraded or low viral load clinical samples in which case human genome normally represent a higher percentage of the sequence data. Compared to long amplicon-based assays, short amplicon-based assays are less affected by RNA degradation and more efficient in generating viral PCR fragments rather than non-specific human fragments as they have shorter lengths of amplicons generated. There is clearly a trade-off between the genome coverage and sensitivity when choosing between short and long amplicon-based assays. Therefore, it has been suggested that long amplicon-based assay could be used as the first round of viral genome amplification and any failed samples could then be amplified with a short amplicon-based assay (26).

From the publicly available RSV full genomes, most have been sequenced with amplicon-based approaches with all ten genes completely obtained but many are lacking a small part of the 155-nt extragenic trailer region at 5′ end of the RSV genome. In addition, most of the RSV-A genomes deposited in NCBI and GISAID databases lacked a large portion of 44-nt extragenic leader region at the 3′ end of the genome (33). Both leader and trailer regions are untranslated, but they were found to be associated with viral replication (34). Notably, these regions are important for designing primers for amplicon-based RSV WGS and using reference sequences missing these nucleotides may account for the failure of both tested short amplicon-based RSV WGS to generate the full *NS1* gene of RSV-A samples.

Mutations were observed in the binding sites of Palivizumab and Nirsevimab in RSV-A and RSV-B we sequenced here, but none of them were associated with increased resistance. Samples were collected in Australia in late 2023 when palivizumab was recommended in the Australian Immunisation Handbook but uptake was not audited and was likely limited, and before the introduction of Nirsevimab. An RSV-A mutation N276S, adjacent to the Palivizumab binding site, was observed in 31 out of 84 the sequences studied. This mutation has been extensively reported and characterized (23). It did not change the neutralization potency of Palivizumab, but that this mutation was associated with selective pressure of Palivizumab and promoted selection of a secondary mutation K272E which become complete resistance to Palivizumab (35). A recent study looking for Nirsevimab escape variants in infants in France during 2023-24 season highlighted the importance of RSV genomic surveillance. In two infants who received one dose of Nirsevimab before their RSV infection, WGS revealed two different mutations in the RSV-B *F* gene (F: N208D or F: I64M + K65R combination), resulting in high levels of Nirsevimab resistance in a fusion-inhibition assay (36). The ONT NGS workflow described here is a powerful tool that can be used for screening of both vaccine and monoclonal breakthrough cases and for general RSV surveillance and evolution studies.

Overall, a rapid and sensitive ONT NGS workflow was established for RSV whole-genome sequencing of clinical respiratory samples based on a long PCR amplicon assay (Supplementary Figure 5). It can also be easily adapted beyond RSV genomic sequencing to include other viruses and pathogens present in clinical respiratory samples such as influenza.

## Author Contributions

Xiaomin Dong: conceptualization, investigation, methodology, data curation, validation, formal analysis, writing – original draft preparation. Steven Edwards: investigation and validation. Yi-Mo Deng: conceptualization, supervision, Writing – Review & Editing. Clyde Dapat and Arada Hirankitti: data curation, formal analysis, and methodology. Rachel Wordsworth: methodology. Paul Whitney, Rob Baird, Kevin Freeman, and Andrew J Daley: Resources. Andrew J Daley: Writing – Review & Editing Ian G. Barr: conceptualization, funding acquisition, supervision, Writing – Review & Editing.

## Acknowledgments

The WHO Collaborating Centre for Reference and Research on Influenza is supported by the Australian Government Department of Health and Aged Care. We would like to thank Nigel Crawford from Murdoch Children’s Research Institute in Australia and Ammar Aziz from Victorian Infectious Diseases Reference Laboratory in the Royal Melbourne Hospital for their advice and resources.

## Data Availability Statement

Sequences generated were deposited into GISAID with accession numbers in supplementary information.

## References

1. Lozano R, Naghavi M, Foreman K, Lim S, Shibuya K, Aboyans V, et al. Global and regional mortality from 235 causes of death for 20 age groups in 1990 and 2010: a systematic analysis for the Global Burden of Disease Study 2010. Lancet. 2012;380(9859):2095–128.

2. Shi T, McAllister DA, O’Brien KL, Simoes EAF, Madhi SA, Gessner BD, et al. Global, regional, and national disease burden estimates of acute lower respiratory infections due to respiratory syncytial virus in young children in 2015: a systematic review and modelling study. Lancet. 2017;390(10098):946–58.

3. Wilkins D, Yuan Y, Chang Y, Aksyuk AA, Nunez BS, Wahlby-Hamren U, et al. Durability of neutralizing RSV antibodies following nirsevimab administration and elicitation of the natural immune response to RSV infection in infants. Nat Med. 2023;29(5):1172–9.

4. Ison MG, Papi A, Athan E, Feldman RG, Langley JM, Lee DG, et al. Efficacy and Safety of Respiratory Syncytial Virus (RSV) Prefusion F Protein Vaccine (RSVPreF3 OA) in Older Adults Over 2 RSV Seasons. Clin Infect Dis. 2024;78(6):1732–44.

5. Mazur NI, Terstappen J, Baral R, Bardaji A, Beutels P, Buchholz UJ, et al. Respiratory syncytial virus prevention within reach: the vaccine and monoclonal antibody landscape. Lancet Infect Dis. 2023;23(1):e2–e21.

6. Melgar M, Britton A, Roper LE, Talbot HK, Long SS, Kotton CN, et al. Use of Respiratory Syncytial Virus Vaccines in Older Adults: Recommendations of the Advisory Committee on Immunization Practices - United States, 2023. MMWR Morb Mortal Wkly Rep. 2023;72(29):793–801.

7. Walsh EE, Perez Marc G, Zareba AM, Falsey AR, Jiang Q, Patton M, et al. Efficacy and Safety of a Bivalent RSV Prefusion F Vaccine in Older Adults. N Engl J Med. 2023;388(16):1465–77.

8. Papi A, Ison MG, Langley JM, Lee DG, Leroux-Roels I, Martinon-Torres F, et al. Respiratory Syncytial Virus Prefusion F Protein Vaccine in Older Adults. N Engl J Med. 2023;388(7):595–608.

9. Efstathiou C, Abidi SH, Harker J, Stevenson NJ. Revisiting respiratory syncytial virus’s interaction with host immunity, towards novel therapeutics. Cell Mol Life Sci. 2020;77(24):5045–58.

10. Liang L, Wang B, Zhang Q, Zhang S, Zhang S. Antibody drugs targeting SARS-CoV-2: Time for a rethink? Biomed Pharmacother. 2024;176:116900.

11. Simoes EAF, Forleo-Neto E, Geba GP, Kamal M, Yang F, Cicirello H, et al. Suptavumab for the Prevention of Medically Attended Respiratory Syncytial Virus Infection in Preterm Infants. Clin Infect Dis. 2021;73(11):e4400–e8.

12. Harvey WT, Carabelli AM, Jackson B, Gupta RK, Thomson EC, Harrison EM, et al. SARS-CoV-2 variants, spike mutations and immune escape. Nat Rev Microbiol. 2021;19(7):409–24.

13. Quick J, Grubaugh ND, Pullan ST, Claro IM, Smith AD, Gangavarapu K, et al. Multiplex PCR method for MinION and Illumina sequencing of Zika and other virus genomes directly from clinical samples. Nat Protoc. 2017;12(6):1261–76.

14. Lagerborg KA, Normandin E, Bauer MR, Adams G, Figueroa K, Loreth C, et al. DNA spike-ins enable confident interpretation of SARS-CoV-2 genomic data from amplicon-based sequencing. bioRxiv. 2021.

15. Grubaugh ND, Gangavarapu K, Quick J, Matteson NL, De Jesus JG, Main BJ, et al. An amplicon-based sequencing framework for accurately measuring intrahost virus diversity using PrimalSeq and iVar. Genome Biol. 2019;20(1):8.

16. Tyson JR, James P, Stoddart D, Sparks N, Wickenhagen A, Hall G, et al. Improvements to the ARTIC multiplex PCR method for SARS-CoV-2 genome sequencing using nanopore. bioRxiv. 2020.

17. Ciuffreda L, Rodriguez-Perez H, Flores C. Nanopore sequencing and its application to the study of microbial communities. Comput Struct Biotechnol J. 2021;19:1497–511.

18. Dong X, Deng YM, Aziz A, Whitney P, Clark J, Harris P, et al. A simplified, amplicon-based method for whole genome sequencing of human respiratory syncytial viruses. J Clin Virol. 2023;161:105423.

19. Wang L, Piedra PA, Avadhanula V, Durigon EL, Machablishvili A, Lopez MR, et al. Duplex real-time RT-PCR assay for detection and subgroup-specific identification of human respiratory syncytial virus. J Virol Methods. 2019;271:113676.

20. Evans D, Kunerth H, Mumm E, Namugenyi S, Plumb M, Bistodeau S, et al. Prospective genomic surveillance and characterization of respiratory syncytial virus in Minnesota, USA – July 2023 to February 2024. medRxiv. 2024:2024.07.10.24310215.

21. Talts T, Mosscrop LG, Williams D, Tregoning JS, Paulo W, Kohli A, et al. Robust and sensitive amplicon-based whole-genome sequencing assay of respiratory syncytial virus subtype A and B. Microbiol Spectr. 2024;12(4):e0306723.

22. Shepard SS, Meno S, Bahl J, Wilson MM, Barnes J, Neuhaus E. Viral deep sequencing needs an adaptive approach: IRMA, the iterative refinement meta-assembler. BMC Genomics. 2016;17(1):708.

23. Hashimoto K, Hosoya M. Neutralizing epitopes of RSV and palivizumab resistance in Japan. Fukushima J Med Sci. 2017;63(3):127–34.

24. Ahani B, Tuffy KM, Aksyuk AA, Wilkins D, Abram ME, Dagan R, et al. Molecular and phenotypic characteristics of RSV infections in infants during two nirsevimab randomized clinical trials. Nat Commun. 2023;14(1):4347.

25. Ribarska T, Bjornstad PM, Sundaram AYM, Gilfillan GD. Optimization of enzymatic fragmentation is crucial to maximize genome coverage: a comparison of library preparation methods for Illumina sequencing. BMC Genomics. 2022;23(1):92.

26. Liu H, Li J, Lin Y, Bo X, Song H, Li K, et al. Assessment of two-pool multiplex long-amplicon nanopore sequencing of SARS-CoV-2. J Med Virol. 2022;94(1):327–34.

27. Kandel S, Hartzell SL, Ingold AK, Turner GA, Kennedy JL, Ussery DW. Genomic surveillance of SARS-CoV-2 using long-range PCR primers. Front Microbiol. 2024;15:1272972.

28. Brejova B, Borsova K, Hodorova V, Cabanova V, Gafurov A, Fricova D, et al. Nanopore sequencing of SARS-CoV-2: Comparison of short and long PCR-tiling amplicon protocols. PLoS One. 2021;16(10):e0259277.

29. Lam C, Johnson-Mackinnon J, Basile K, Fong W, Suster CJ, Gall M, et al. A laboratory framework for ongoing optimization of amplification-based genomic surveillance programs. Microbiol Spectr. 2023;11(6):e0220223.

30. Itokawa K, Sekizuka T, Hashino M, Tanaka R, Kuroda M. Disentangling primer interactions improves SARS-CoV-2 genome sequencing by multiplex tiling PCR. PLoS One. 2020;15(9):e0239403.

31. Davina-Nunez C, Perez-Castro S, Cabrera-Alvargonzalez JJ, Montano-Barrientos J, Godoy-Diz M, Regueiro B. The Modification of the Illumina((R)) CovidSeq Workflow for RSV Genomic Surveillance: The Genetic Variability of RSV during the 2022-2023 Season in Northwest Spain. Int J Mol Sci. 2023;24(22).

32. Freed NE, Vlkova M, Faisal MB, Silander OK. Rapid and inexpensive whole-genome sequencing of SARS-CoV-2 using 1200 bp tiled amplicons and Oxford Nanopore Rapid Barcoding. Biol Methods Protoc. 2020;5(1):bpaa014.

33. Collins PL, Fearns R, Graham BS. Respiratory syncytial virus: virology, reverse genetics, and pathogenesis of disease. Curr Top Microbiol Immunol. 2013;372:3–38.

34. Hanley LL, McGivern DR, Teng MN, Djang R, Collins PL, Fearns R. Roles of the respiratory syncytial virus trailer region: effects of mutations on genome production and stress granule formation. Virology. 2010;406(2):241–52.

35. Zhu Q, Patel NK, McAuliffe JM, Zhu W, Wachter L, McCarthy MP, et al. Natural polymorphisms and resistance-associated mutations in the fusion protein of respiratory syncytial virus (RSV): effects on RSV susceptibility to palivizumab. J Infect Dis. 2012;205(4):635–8.

36. Fourati S, Reslan A, Bourret J, Casalegno JS, Rahou Y, Chollet L, et al. Genotypic and phenotypic characterisation of respiratory syncytial virus after nirsevimab breakthrough infections: a large, multicentre, observational, real-world study. Lancet Infect Dis. 2024.

