## Supplemental material 1 for "An improved rapid and sensitive long amplicon method for nanopore-based RSV whole genome sequencing"

### Supplementary Tables

Supplementary Table 1. The information for primers used in the modified one-step multiplex RT-PCR, note that changes have been made to PCR primer concentration, thermal conditions, and primers of amplifying fragment 3 and 6 compared to our previous publication (PMID: 36934591

| Multiplex Tube | Primer Name | Primer Sequence (5'- 3')[b] | Position of first 5' base[a] | Amplicon length (bp) | Final concentration (μM) |
|---|---|---|---|---|---|
| Tube 1 | 1-1F | ACGCGAAAAAATGCGTACTACAAAC | 1 | | 0.16 |
| | 1-1R | CTG**M**ACCATAGGCATTCATAAACA | 1898 | 1898 (fragment 1) | 0.16 |
| | 1-3F_3 | GCATCACT**W**ACAATATGGGT**K**CC | 2926 | | 0.55 |
| | 1-3R_6 | CACTTTTGAT**Y**TTGTTCACTTC**Y**CC | 6304 | 2876 (fragment 3) | 0.55 |
| | 1-5F | TGATGCATCAATATCTCAAGTCA | 7181 | | 0.6 |
| | 1-5R | G**R**CCTAT**D**CCTGCATACTC | 11116 | 3936 (fragment 5) | 0.6 |
| Tube 2 | 2-2F | ATGGGAGA**R**GT**R**GCTCCAGAATA | 1562 | | 0.25 |
| | 2-2R | CGTGTAGCTGT**R**TG**Y**TTCCAA | 4004 | 2443 (fragment 2) | 0.25 |
| | 2-4F | AGCAAATT**Y**TGGCC**Y**TA**Y**TTTAC | 4334 | | 0.6 |
| | 2-4R1 | CTCATAGCAACACATGCTGATTG | 7967 | 3634 (fragment 4) | 0.3 |
| | 2-4R2 | GAGTTTGCTCATGGCAACACAT | 7974 | 3641 (fragment 4) | 0.3 |
| | 2-6F | TGGACCAT**W**GAAGC**Y**ATATCA | 10912 | | 0.44 |
| | 2-6R | AGTGTCAAAAACTAAT**R**TCTCGT | 15265 | 4354 (fragment 6) | 0.44 |
| | 2-6R2 | ACGAGAAAAAAAGTGTCAAAAACTAAT | 15272 | 4365 (fragment 6) | 0.44 |

[a]Primer nucleotide numbering was based on a human RSV-B strain (GISAID accession numbers 2584506)

[b]Degenerate bases are highlighted in bold.

Supplementary Table 2. Component for multiplex RT-PCR

| Component for multiplex PCR mix 1 | Volumes (µL) per reaction |
|---:|---:|
| 1-1F | 0.2 |
| 1-1R | 0.2 |
| 1-3F_3 | 0.69 |
| 1-3R_6 | 0.69 |
| 1-5F | 0.75 |
| 1-5R | 0.75 |
| 2X Platinum SuperFi RT-PCR Master Mix | 12.5 |
| SuperScript IV RT Mix | 0.25 |
| template RNA | 8.97 |

| Component for multiplex PCR mix 2 | Volumes (µL) per reaction |
|---:|---:|
| 2-2F | 0.31 |
| 2-2R | 0.31 |
| 2-4F | 0.75 |
| 2-4R1 | 0.38 |
| 2-4R2 | 0.38 |
| 2-6F | 0.55 |
| 2-6R | 0.55 |
| 2-6R2 | 0.55 |
| 2X Platinum SuperFi RT-PCR Master Mix | 12.5 |
| SuperScript IV RT Mix | 0.25 |
| template RNA | 8.47 |

Supplementary Table 3. Comparison of sequencing samples with insufficient amplicons by using rapid barcoding (RBK) and rapid PCR barcoding (RPB) kits.

NGS results were characterised into four groups including Full-length (RSV whole genome sequence obtained), G (Full G sequence only), F (Full F sequence only), GF (Full F and G sequence obtained) and other (no full G and F sequences obtained).

| Sample ID | Ct Value | NGS type | NGS RPB | NGS RBK |
|---|---|---|---|---|
| **90007809** | 29 | A | GF | Full-length |
| **90007814** | 30 | B | Other | Full-length |
| **90007817** | 27 | B | G | Full-length |
| **90007819** | 30 | B | G | Other |
| **90008162** | 25 | B | G | Full-length |
| **90008175** | 24 | A | GF | Full-length |
| **90008179** | 26 | A | GF | Full-length |
| **90008178** | 28 | A | G | Full-length |
| **90008183** | 23 | B | GF | Full-length |
| **90008191** | 27 | A | GF | Full-length |
| **90008187** | 27 | A | GF | Full-length |
| **90008196** | 27 | A | GF | Full-length |
| **90008203** | 33 | A | G | Other |
| **90008227** | 28 | B | G | G |
| **90008229** | 32 | A | G | G |
| **90008225** | 27 | A | GF | Full-length |
| **90008228** | 28 | A | G | Full-length |
| **90008224** | 30 | A | GF | G |
| **90008231** | 26 | A | G | Full-length |
| **90008380** | 35 | A | Other | Other |

### Supplementary Figure

A

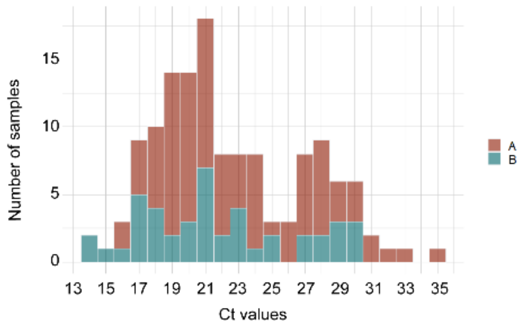


B

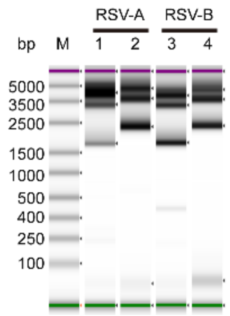


Supplementary Figure 1. The performance of modified RSV mRT-PCR for generating RSV genomic amplicons in 135 clinical samples.

(A) Histograms of the Ct value distribution for clinical samples with two RSV subgroups, RSV-A in dark red and RSV-B in dark green. (B) Tapestation images of RSV amplicons generated from one RSV-A and one RSV-B positive clinical samples.

A

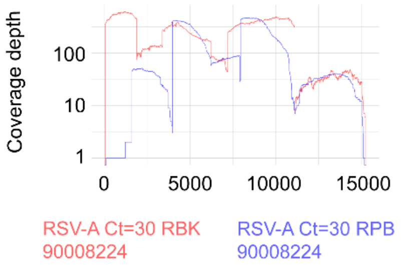


B

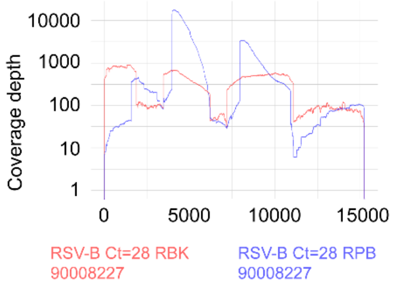


Supplementary Figure 2. RSV NGS achieved by ONT rapid barcoding (RBK) and rapid PCR barcoding (RPB) for samples with insufficient amplicons for NGS library preparation.

Coverage depth of sequenced representative RSV-A (A) and RSV-B (B) in genomic position with RBK (red) and RPB (blue) for NGS library preparation

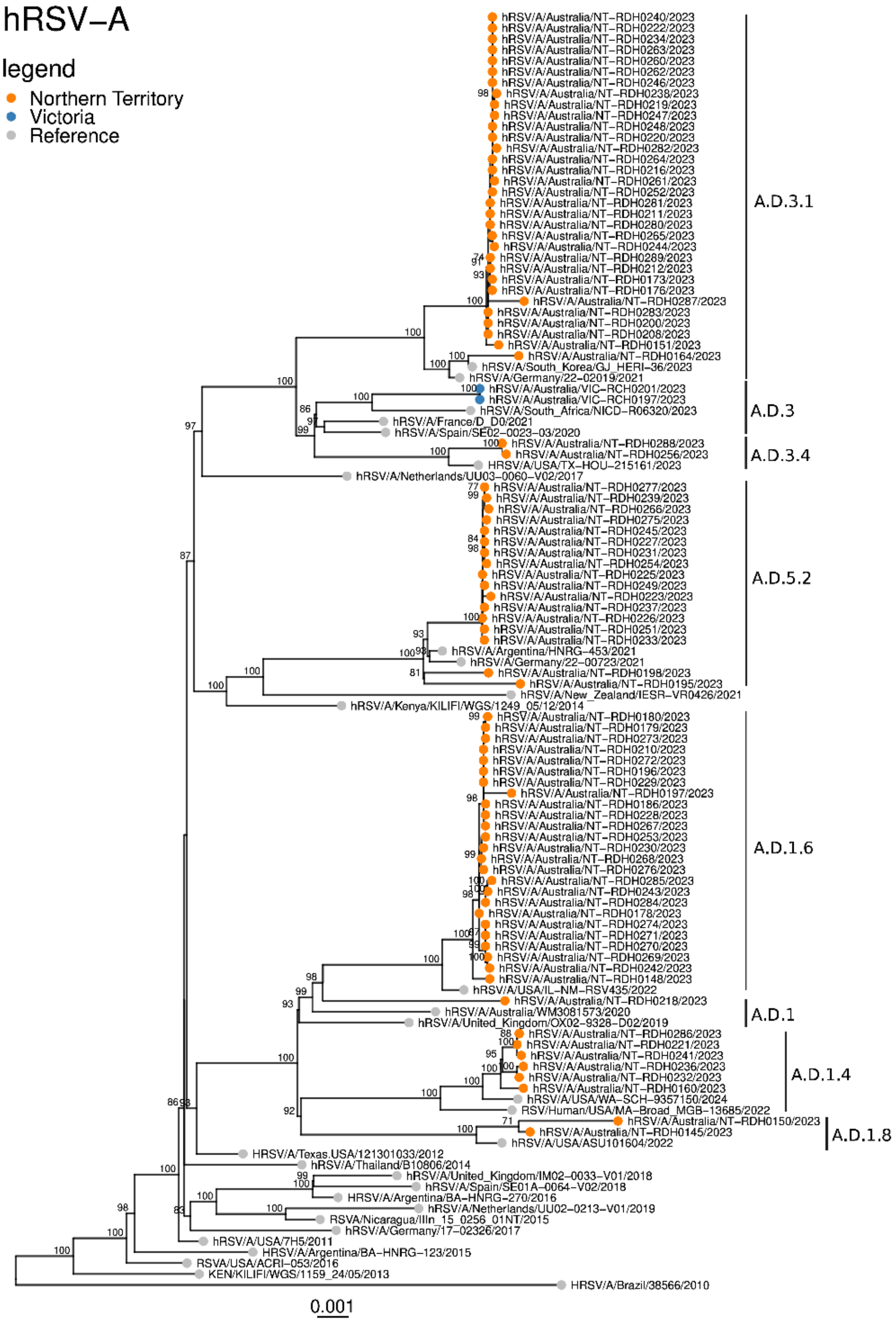


Supplementary Figure 3. Phylogenetic analysis of full-length of RSV-A

Phylogenetic analysis of full-length of RSV-A generated using the ONT workflow described in this method. RSV full-length genome sequences generated using this method were analyzed phylogenetically and showed that the RSV-A genomes clustered into 3 main clades.

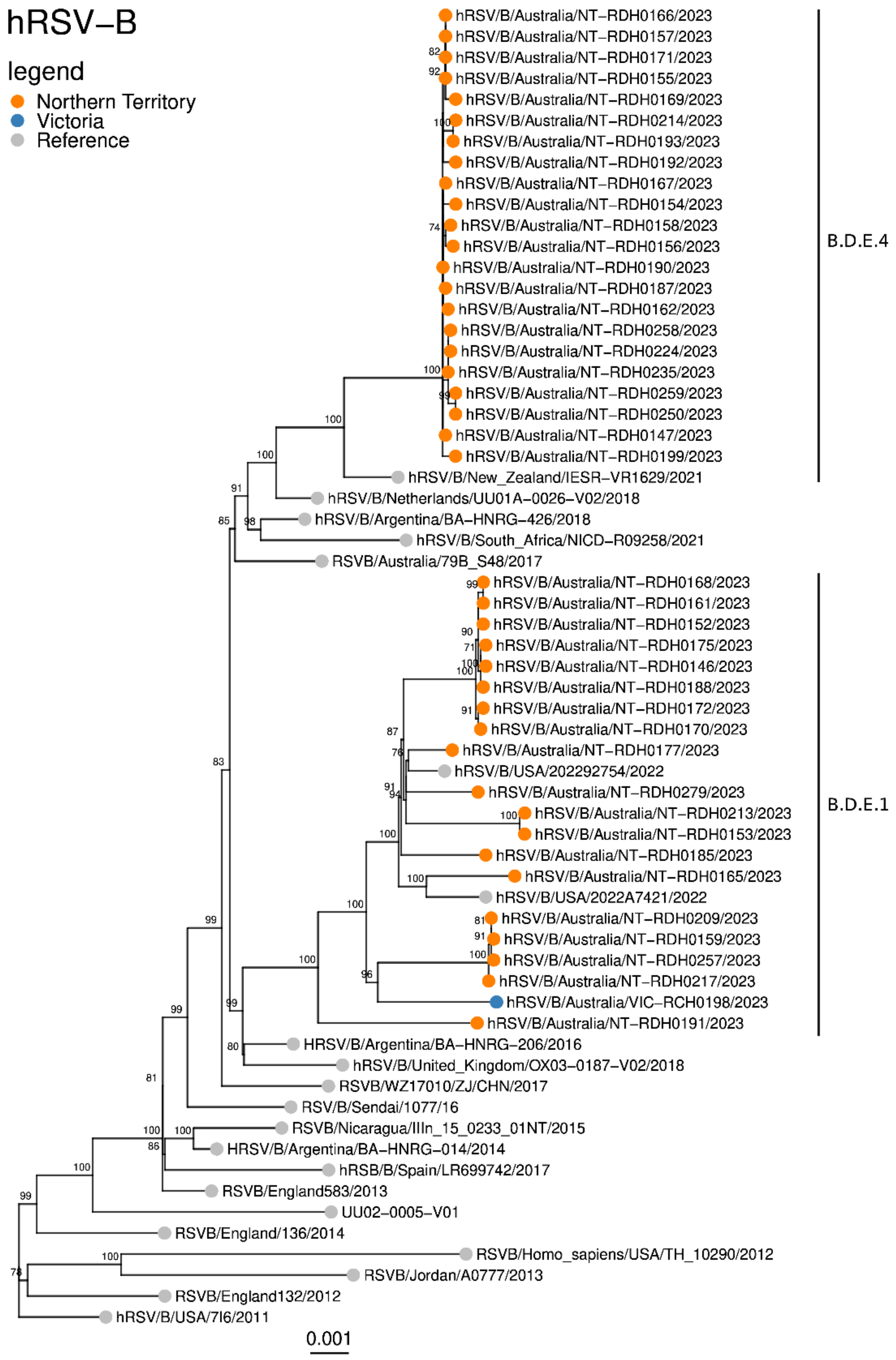


Supplementary Figure 4. Phylogenetic analysis of full-length of RSV-B

Phylogenetic analysis of full-length of RSV-B generated using the ONT workflow described in this method. RSV full-length genome sequences generated using this method were analyzed phylogenetically and showed that the RSV-B genomes clustered into 2 main clades.

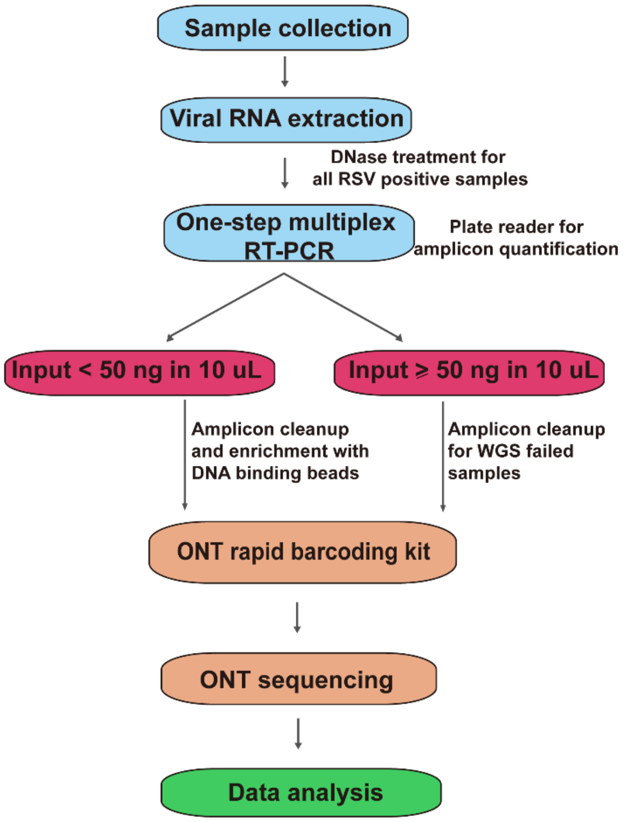


Supplementary Figure 5. Schematic diagram of ONT NGS workflow for RSV WGS based on a long PCR amplicon assay.
